# Post-Infection Entry Mechanism of Ricin A Chain-Pokeweed Antiviral Proteins (RTA-PAPs) Chimeras is Mediated by Viroporins

**DOI:** 10.1101/2021.06.17.448882

**Authors:** Yasser Hassan

## Abstract

The limitations of virus-specific antiviral drugs became apparent during the current COVID-19 pandemic. The search for broad range antiviral proteins of a new kind to answer current and future pandemics has become an even more pressing matter. Here, the author further describes the expected anti-SARS-CoV-2 mechanisms of a novel broad range antiviral chimeric protein constructed between ricin A chain and pokeweed antiviral proteins. The latest in protein-ligand docking software were used to determine binding affinity of RTA-PAPs to SARS-CoV-2 frameshift stimulation element and elucidate the preferential post-infection entry mechanisms of RTA-PAPs into virus infected cells over non-infected ones, by doing a comparative analysis between in vitro and in silico results on numerous viruses. The results obtained strongly suggest that the post-infection preferential entry of RTA-PAPs into infected cells is mediated by the presence of viroporins integrated into the host cell membrane. The discovery of this mechanism revealed RTA-PAPs, and proteins like them, to be a new class of broad range antivirals that target with high specificity viroporin producing viruses, and with gain of functions in antiviral activities, post-infection.

## 1. Introduction

Scientists around the globe are actively searching for potent antiviral drugs to treat and cure all kind of diseases caused by various viral infections. The main strategies to develop such antivirals have been focused on small virus-specific molecules that hinder a precise mechanism in the viral cycle, over time, and on neutralizing antibodies, i.e. vaccines or monoclonal antibodies. Those strategies have been successful so far, however, they are not without limitations. Indeed, the small virus-specific molecules lead to mutated forms of the viruses, over time, that are resistant to treatments, and the development of neutralizing antibodies is a lengthy and costly process. In order to develop virus-specific vaccines or monoclonal antibodies, an average of five to ten years is required [1]. Additionally, those strategies are reactive as opposed to proactive, i.e. those antivirals are only developed after the viral infections happened.

The current COVID-19 pandemic, caused by the severe acute respiratory syndrome coronavirus 2 (SARS-CoV-2) infection, brought to light those limitations. Indeed, there are still no efficient antivirals against SARS-CoV-2, and the existing vaccines, while efficient, need to be renewed due to the emergence of new variants of the virus, capable of evading the immune system. The need for broad range antiviral drugs capable to treat numerous variants of the same virus and viruses of the same family became apparent with the ongoing COVID-19 pandemic.

There are many potential strategies to develop broad range antivirals, and one of those strategies of interest is to look into plants defensive mechanisms. Plants have evolved numerous proteins capable of fighting off not only viral infections but also bacterial and fungal ones, using direct and indirect mechanisms. Ribosome inactivating proteins (RIPs) are such proteins, and can irreversibly inactive protein synthesis, leading to cell death, upon infection by a foreign pathogen [2]. Interestingly, certain RIPs have been found to have very potent direct activities against various animal and human viruses by inhibiting viral entry, hindering viral cell machinery and depurinating viral RNA/DNA. Furthermore, certain RIPs gain entry into infected cells preferentially over non-infected ones by an unclear mechanism. For those reasons, RIPs have been the subject of extensive investigations in order to circumvent the limitations of virus-specific antivirals.

Among the most studied RIPs over the last forty years are pokeweed antiviral proteins (PAPs) and ricin. PAPs and ricin were found to be very potent against numerous human viruses such as Japanese encephalitis virus, human immunodeficiency virus-1 (HIV-1), human T-cell leukemia virus-1, herpes simplex virus, influenza, hepatitis B virus (HBV), and poliovirus in vitro and in clinical trials [3]. However, RIPs suffer dose limiting toxicity and unfavorable pharmacokinetics (PK) profiles when administered alone. RIPs are mostly used today as immunotoxins or conjugated toxins. Nonetheless, the author designed chimeras constructed with PAPs and ricin-A chain (RTA) (RTA-PAPs) with gain of function in protein synthesis inhibition, and in antiviral activity in vitro and in silico against numerous human viruses, such as HBV and SARS-CoV-2, at subtoxic dosages with a better PK profile than either moieties alone [3–4].

Given the importance of developing broad range antivirals, the author hypothesized that based on the previous results published [3–5], and using state of the arts in silico tools, it would be possible to elucidate RTA-PAPs preferential entry mechanism into infected cells over non-infected ones, and further our understanding of the expected anti-SARS-CoV-2 mechanisms discussed previously [4]. Here, the author reports the latest results regarding the novel binding mechanisms of RTA-PAPs against the frameshift stimulation element (FSE) of SARS-CoV-2 genome, required for balanced expression of essential viral proteins [6], the novel associated potential depurination potential, and the logically deduced preferential entry mechanism into infected cells over non-infected ones based on all the accumulated in vitro, and newly generated in silico results.

## 2. Results

The three-dimensional (3D) structure of SARS-CoV-2 FSE was made available recently [6], and was acquired from the Research Collaboratory for Structural Bioinformatics (RCSB) website (https://www.rcsb.org/). The 3D structures of RTA, PAP1 and PAPS1 were also retrieved from RCSB. The predicted 3D structures of the fusion protein between RTA and pokeweed antiviral protein from seed isoform 1 (PAPS1) (RTA-PAPS1), and the chimera between RTA mutant (RTAM) and pokeweed antiviral protein isolated from leaf isoform 1 (PAP1) (RTAM-PAP1) were obtained as previously described [3]. An additional stable conformation of RTAM-PAP1 (RTAM-PAP1 2^nd^) was discovered in the process of double checking all the constructs, and was generated using the same methodology used for RTAM-PAP1 described in our previous publications [3–4]. The new conformation is more similar in structure to RTA-PAPS1 than RTAM-PAP1, in the orientation of the PAP1 moiety (figure 1.a-c). Furthermore, an analysis of RTAM-PAP1 2^nd^ by I-TASSER showed that this particular conformation might have the ability to bind and depurinate Uracil (Appendix A. Figure 1.), on top of the usual Adenine and occasional Guanine [3]. A structure comparison was done using MATRAS to confirm those results, and RTAM-PAP1 2^nd^ was found to be very similar, in structure, to Uracil-DNA-Glycosylase at the uracil binding cleft and catalytic site (Appendix A. Figure 2.) The sequence identity was of 5.3% while the secondary structure similarity was of 65.8% between amino acids 367-410 of RTAM-PAP1 2^nd^ and 154-191 of Uracil-DNA-Glycosylase (UDG).

**Figure 1.**
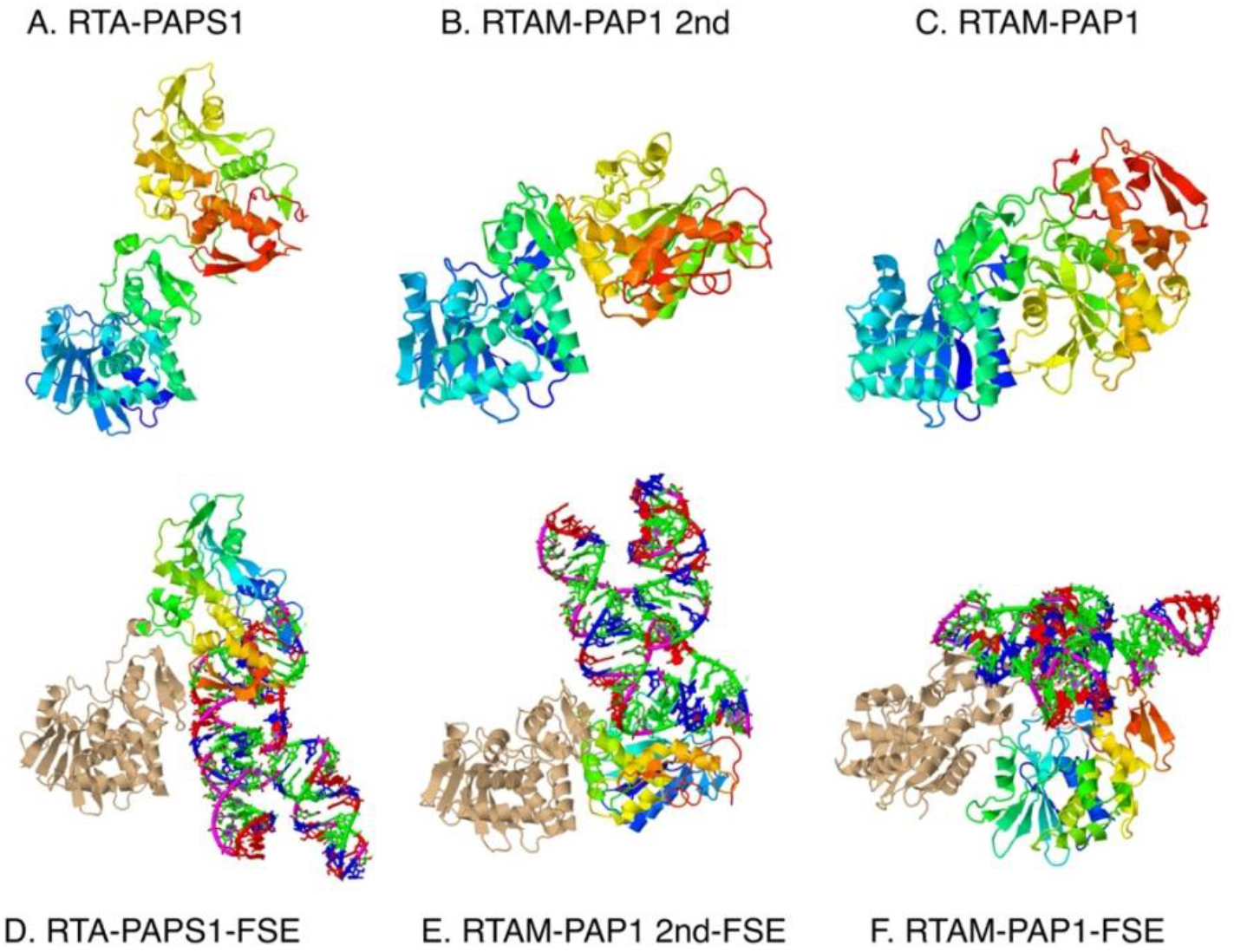
The predicted 3D models generated by I-TASSER of A) RTA-PAPS1, B) RTAM-PAP1 2nd and C) RTAM-PAP1. The conformation of RTAM-PAP1 2nd more closely resembles that of RTA-PAPS1. N to C terminal in blue to red, with RTA(M) moieties being on the left, mostly blue and green. The top 3D models of the different RTA-PAPs chimeras in complex with FSE generated by ZDOCK in carton format are shown in D) for RTA-PAPS1-FSE, E) RTAM-PAP1 2nd -FSE and F) RTAM-PAP1-FSE. The RTA(M) moieties are colored in beige, the PAPs moieties in blue to red, N to C, and the FSE in RNA colors (green, red, blue and purple). All the models were viewed in Jena3D. FSE is “neatly” in the main catalytic cleft for RTAM-PAP1 2nd-FSE supported by RTA.

A hybrid algorithm of template-based and template-free docking was performed for the three compounds, RTA-PAPS1, RTAM-PAP1 and RTAM-PAP1 2^nd^, against SARS-CoV-2 FSE (Protein-RNA docking) using the HDOCK webserver [7]. The results (table 1) clearly show the gain of function on binding energies for the three chimeras in comparison with the moieties alone. All of the fusion proteins bind SARS-CoV-2 FSE with both moieties, but RTAM-PAP1 2^nd^ was found to have the highest binding affinity (lowest binding energy) for FSE, which was achieved within the catalytic cleft of the PAP1 moiety and on the putative Uracil biding site.

**Table 1.** The binding energies in kcal/mol for the models generated by HDOCK for each compound in complex with the RNA frameshift site (FSE). Top 1 is the model with the lowest binding energy (highest binding affinity) and the top 10 is the 10th model with the lowest binding energy. The lowest energy for the top 1 and top 10 models is in bold.

| SARS-CoV-2 |
| --- |
| RNA Frameshift Site |

|  | Top 1 | Top 10 |
| --- | --- | --- |
| Compounds |  |  |
| RTA-PAPS1 | -354 | -319 |
| RTAM-PAP1 | -375 | **-330** |
| RTAM-PAP1 2<sup>nd</sup> | **-426** | -318 |
| RTA | -345 | -297 |
| PAP1 | -311 | -287 |
| PAPS1 | -326 | -290 |

In order to confirm the probability of purine or pyrimidine removal, a protein-ligand docking was run using Z-DOCK for the three chimeras against SARS-CoV-2 FSE (figure 1.d-f). The binding conformation of RTAM-PAP1 2^nd^ to SARS-CoV-2 FSE was again found to lie just within the catalytic cleft of PAP1 moiety. This conformation was found to be extremely stable, with the lowest binding energy (−426 kcal/mol) among the three fusion proteins, and is associated with depurinating activities [3–4].

The different chimeras have comparable binding energies, but with a noticeable difference for RTAM-PAP1 2^nd^. This doesn’t come as a surprise since it is a known fact that protein 3D structure determines function, and, thus, any conformational change will incur most of the time a functional change. It is also interesting to note that PAPS1 has a higher binding affinity (−326 kcal/mol) to SARS-CoV-2 FSE than PAP1 (−311 kcal/mol), and, yet, RTA-PAPS1 has the lowest binding affinity (−354 kcal/mol) to SARS-CoV-2 FSE of the chimeras. Given these new results, and the fact that RTA-PAPs were already found to have very high binding affinity towards SARS-CoV-2 key proteins, comparable to that of convalescent COVID-19 patient-origin B38 antibody to SARS-CoV-2 Spike protein [4], it is safe to assume that RTA-PAPs will have both direct and indirect anti-SARS-CoV-2 activities.

However, one question remains, will RTA-PAPs be able to enter preferentially infected cells over non-infected ones to exert their antiviral activities, and, thus, have a high Therapeutic Index (TI)? The understood mechanism for the preferential entry into infected cells is during the virus adsorption phase, virions modify their conformation upon receptor binding allowing the insertion of virion components into the cellular membrane, inducing changes in membrane permeability. The viral components open pores inducing a membrane potential drop allowing macromolecules to enter through translocation, pushed through the cell membrane by a proton motive force [8].

Given the structure and function similarities between the different RTA-PAPs, a comparative study of in vitro results and in silico results was performed in order to determine common factors to preferential entry into infected cells and antiviral activity. Results of RTA-PAPS1 antiviral activities in vitro were previously published [3, 5], and RTA-PAPS1 was found to have antiviral activity post-infection in cytoprotection assays against five out of nine viruses tested with a TI greater than 1. (table 2).

**Table 2.**
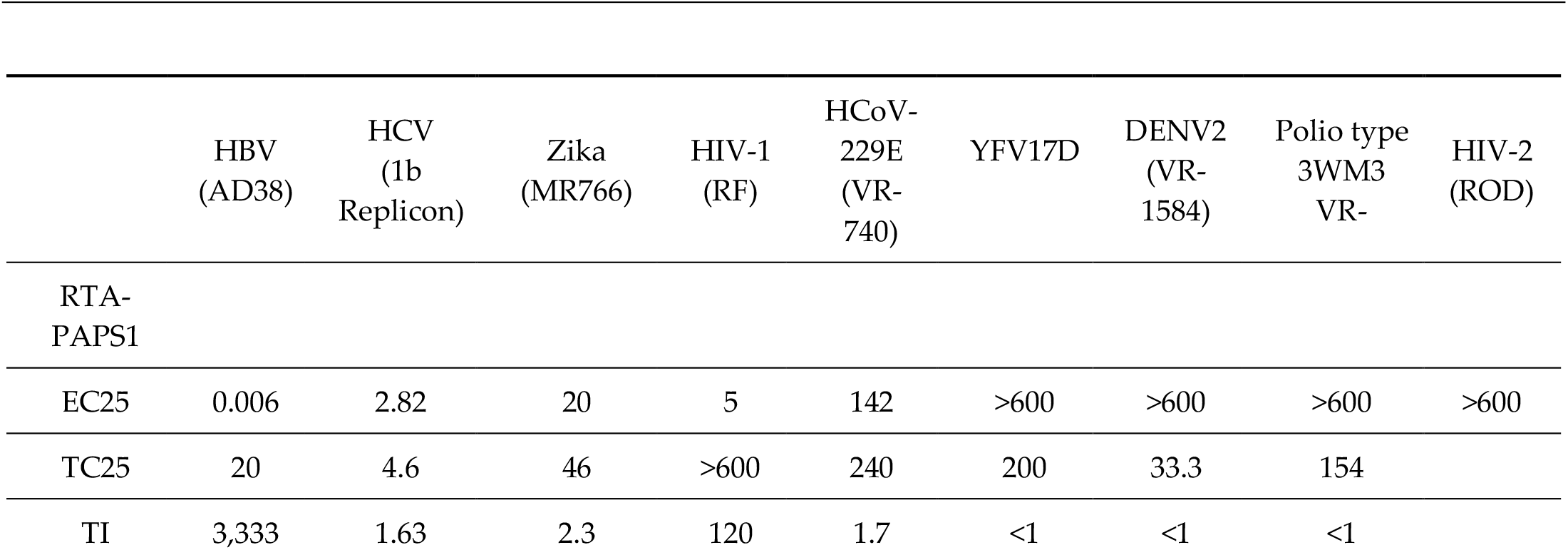
The post-infection concentrations of RTA-PAPS1 needed to inhibit 25% cytopathic effects (EC25) of various viruses on cells compared to amount needed to be toxic to healthy cells causing 25% less growth (TC25) in nanoMolar (nM). The Therapeutic Index (TI) represents the ratio TC25/EC25 and is an indicator of RTA-PAPS1 preferential entry and antiviral activity in infected cells over non-infected cells. A value less than 1 means no antiviral activity before becoming toxic to healthy cells.

The most obvious common structure between those viruses with a strong positive correlation with TI > 1 are viroporins that get integrated into the host cell membrane (*Pearson’s r*(9) = 0.8803, *p* < .01, calculations in Appendix B.). Viroporins are, for the most part, small viral proteins that oligomerize in the membrane of host cells and modify several cellular functions, including Ca^2+^ homeostasis and membrane permeability, i.e. form ion channels. They have been found to be directly linked to viral virulence [9]. Indeed, HBV [10–11], HCV [12], Zika virus [13], HIV-1 [14] and HCoV-229 [15] all express viroporins. On the other hand, HIV-2 [14], YFV17D [16], Poliovirus [17] and DENV2 [18] do not express viroporins, or the strain tested is attenuated with at least one mutation in the viroporin gene (i.e. less virulent strain), or the viroporin does not integrate host cell membrane (i.e. Polioviruses). The YFV-17D, albeit similar to Zika virus, is actually a live attenuated virus used for vaccines with multiple mutations in its viroporin gene among others, for example. Zika MR766 strain does not have mutations in its viroporin gene. These results are very much in line with the current understanding of the preferential entry mechanism, and, actually, confirm the correlation between the presence of viroporins for RTA-PAPS1 to gain preferential entry into infected cells over non-infected cells, post infection.

In order to determine if there is a correlation between viral proteins integrated into host cell membranes during and after viral infection and TI of RTA-PAPS1, a knowledge-based scoring docking prediction was performed for all the compounds against the viral membrane proteins and viroporins of HBV, HIV-1, HCoV-229 and SARS-CoV-2 (table 3) using CoDockPP global docking.

**Table 3.**
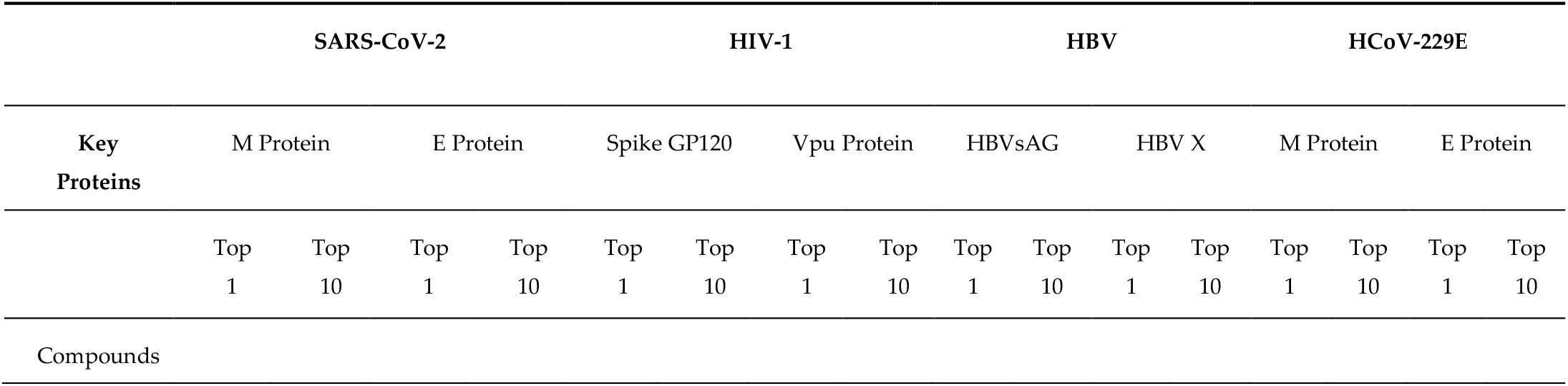
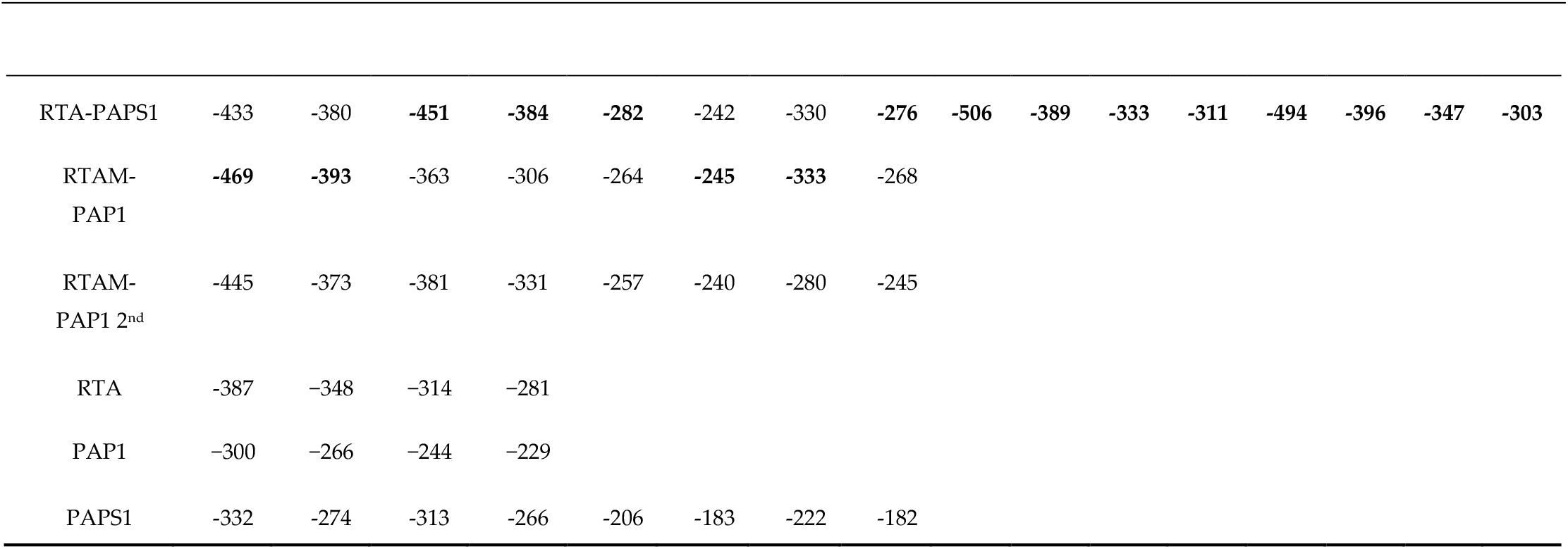
The binding energies in kcal/mol for the models generated by CoDockPP for each compound in complex with the outer virus main envelope protein and viroporin that integrate host cell membrane. Top 1 is the model with the lowest binding energy (highest binding affinity) and the top 10 is the 10th model with the lowest binding energy. The lowest energy for the top 1 and top 10 models for each complex is in bold.

The results show that all the chimeras have comparable high binding affinity toward all the viroporins and increasing binding affinity toward viral membrane proteins with increasing TI against respective viruses. The only exception is the very high binding affinity of RTA-PAPS1 toward wild type HCoV-229E M and E proteins with an in vitro TI of 1.7. Again, this can be explained by many factors, but the fact that the strain tested in vitro is a less virulent one, i.e. mutations in both the M and E proteins, it is conceivable that RTA-PAPS1 is actually very active against wild type HCoV-229E. Albeit there are not enough data to determine a clear correlation between antiviral activity and binding affinity to envelope proteins, those results make sense. Indeed, the high binding affinity towards viral proteins that get integrated into the cell membrane means that RTA-PAPs spend more time interacting with said proteins and, thus, are more inclined to enter cells infected by those particular viruses. The higher rate of entry into infected cells over non-infected ones by RTA-PAPs should correlate with higher TI. There are, of course, many other factors that come into play, such as type of antiviral activity (i.e. depurination activity of viral RNA, for example). This mechanism should nonetheless shed light on why RTA-PAPS1 was so active against HBV in vitro (TI of 3333), and, also, help explain the preferential entry mechanisms into infected cells over non infected ones of certain type 1 RIPs such as PAPs. Based on the data presented, it would be safe to assume that RTA-PAPs will be very active against SARS-CoV-2, its virulent variants, coronaviruses in general, and, any viroporin producing viruses with key proteins RTA-PAPs have high binding affinity to. Indeed, SARS-CoV-2 FSE is highly conserved across a wide range of coronaviruses, a membrane M and E protein are expressed by most coronaviruses, and any viroporin producing virus, that integrates the host cell membrane, should allow preferential entry of RTA-PAPs into infected cells, opening the door to induction of cell apoptosis at the very least.

## 3. Conclusion

In conclusion, RTA-PAPs are effectively a new class of broad range antiviral proteins that target with high specificity viruses that produce viroporins that get integrated into the host cell membrane. The rate at which RTA-PAPs will enter post-infection preferentially virus infected cells over non-infected ones should be directly dependent on the binding affinity to host cell membrane integrated viral envelop proteins and viroporins. The type of the viroporins might be an important factor in said rate as it might affect the proton motive force required for RTA-PAPs to go through the cell membrane. However, the actual antiviral activity of RTA-PAPs will depend on other factors, such as ability to depurinate viral DNA/RNA efficiently or to induce cell apoptosis in infected cells. It would be interesting to find out if other chimeras constructed with toxins will have similar gain of functions or entirely new ones.

## 4. Materials and Methods

### 4.1 Protein Modeling

#### 4.1.1. Generation of 3D Structures

The predicted molecular 3D structures of RTA-PAPS1 and RTAM-PAP1 were already available from previous work [3–4] and are available in the supplementary files (PDB files S1). The predicted molecular 3D structure of RTAM-PAP1 2^nd^ was generated the same way as RTAM-PAP1 [3] and is available in the supplementary files (PDB file S1) as well. The 3D models for RTA, PAP1 and PAPS1 were retrieved from RCSB (https://www.rcsb.org), in PDB format with the following PDB ID: 4MX5, 1PAG and 1J1S respectively. The 3D models for SARS-CoV-2 M and E proteins were retrieved directly from the CoDockPP site (https://ncov.schanglab.org.cn/). The 3D models for HIV-1 GP120 and Vpu proteins were retrieved from RCSB in PDB format with the following PDB ID: 5F4U and 1VPU respectively. The predicted molecular 3D structures for HBV X, HBVsAG, HCoV-229e M and HCoV-229e E proteins were generated using Phyre2 prediction server [19] only. The 3D structures weren’t refined further as it was not needed for the experiments to have a more accurate prediction to be meaningful, and are available in the supplementary files (PDB file S1). The 3D structure of SARS-CoV-2 FSE was retrieved from RCSB in PDB format with the PDB ID: 6XRZ.

#### 4.1.2. Structure Modeling

The structures of the Protein-Protein bound complexes were generated by CoDockPP using the ambiguous peptide-ligand computations [20–22]. RTAM-PAP1 2^nd^ was compared by superimposition on the available crystallography in RCSB using MATRAS pairwise 3D alignment (http://strcomp.protein.osaka-u.ac.jp/matras/matras_pair.html). The structures of the Protein-RNA bound complexes were generated by a hybrid algorithm of template-based and template-free docking using HDOCK webserver [7]. Additional models of the RTA-PAPS1, RTAM-PAP1, RTAM-PAP1 2^nd^, RTA-PAPS1-FSE, RTAM-PAP1-FSE and RTAM-PAP1 2nd-FSE complexes were generated using ZDOCK [23] without inputting the active residues. All models were viewed using Jena3D (http://jena3d.leibniz-fli.de/).

## Supplementary Materials

### Author Contributions

Y.H. is the sole author.

### Funding

This research received no external funding”.

### Conflicts of Interest

“The author is a shareholder at Ophiuchus Medicine Inc. which owns the patent rights to described RTA-PAPs chimeras.”

## Appendix A

**Figure 1.**
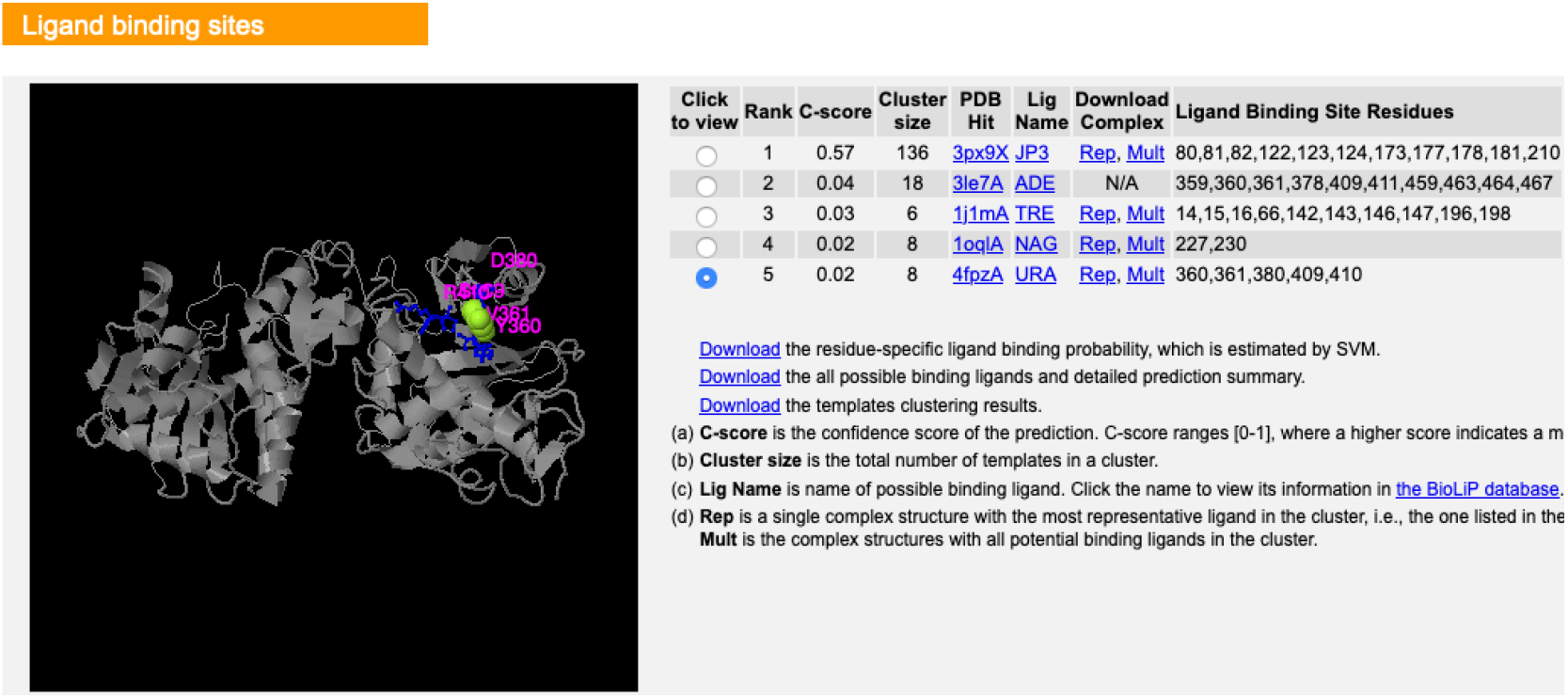
Results from I-TASSER showing that Uracil might be a ligand to RTAM-PAP1 new conformation (shown in black box with RTAM on the left) with binding sites 360,361,380,409 and 410.

**Figure 2.**
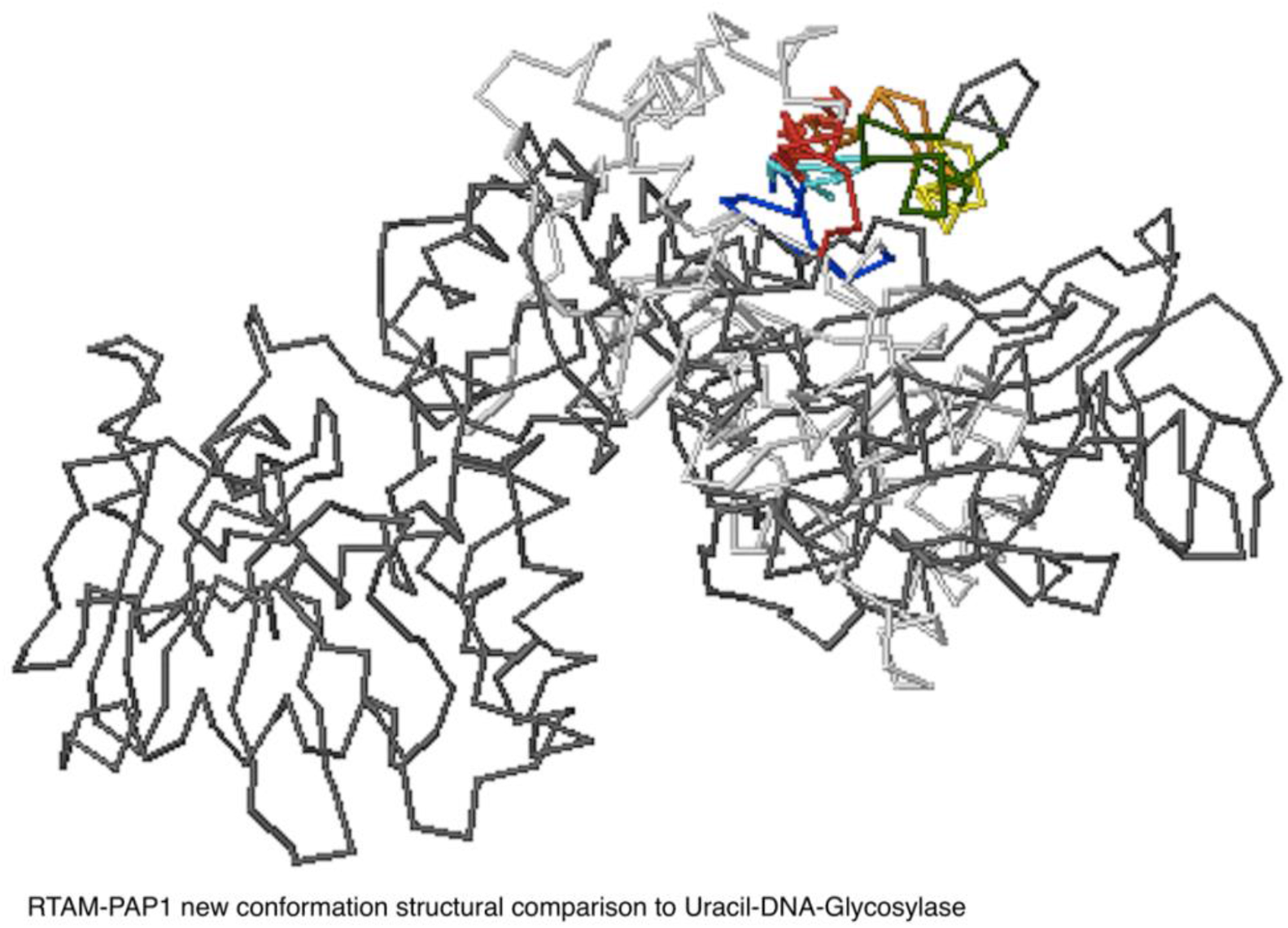
RTAM-PAP1 2nd structural comparison to Uracil-DNA-Glycosylase. RTAM-PAP1 is in black (RTAM on the left side) and UDG is in grey. The structural similarities are in colors.

## Appendix B

Pearson correlation coefficient calculations between presence of viroporin that get integrated into host cell membrane and Therapeutic Index (TI) >1.

*Key*

*X: Presence of viroporin that get integrated into host cell membrane in virus (Yes =1, No =0, presence of mutated viroporin (attenuated) = 0*.*5)*

*Y: TI of RTA-PAPS1 against virus (TI > 1 = 1, TI < 1 = 0)*

*M*_*x*_: *Mean of X Values*

*M*_*y*_: *Mean of Y Values*

*X - M*_*x*_ *& Y - M*_*y*_: *Deviation scores*

*(X - M*_*x*_)^*2*^ *& (Y - M*_*y*_)^*2*^: *Deviation Squared*

*(X - M*_*x*_*)(Y - M*_*y*_*): Product of Deviation Scores*

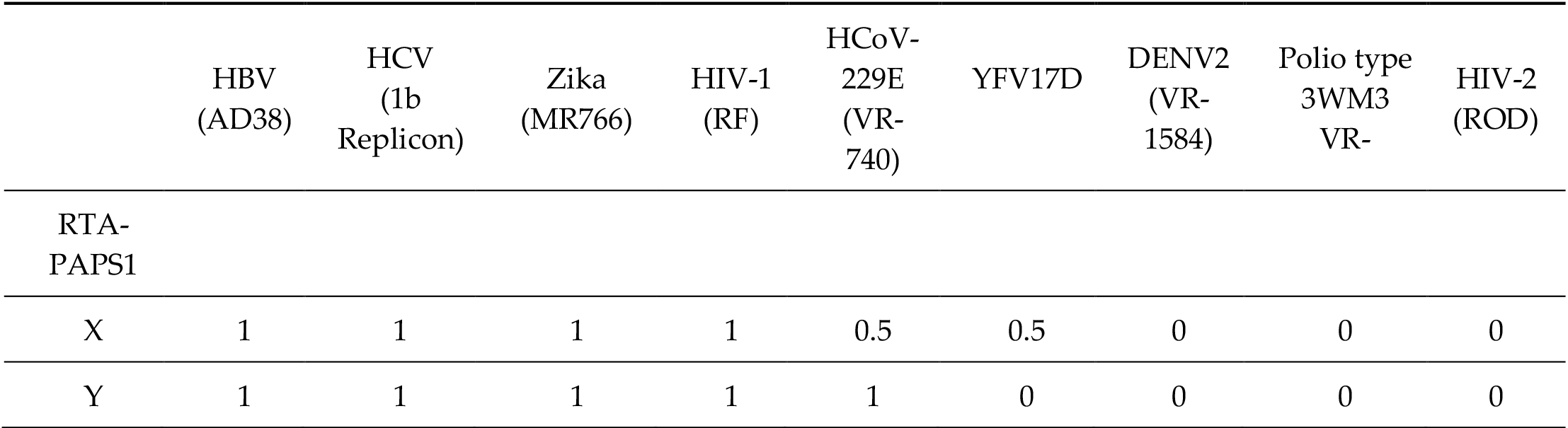

*Result Details & Calculation*

*X Values*

∑ = 5

Mean = 0.556

∑ (X - M_x_)^2^ = SS_x_ = 1.722

*Y Values*

∑ = 5

Mean = 0.556

∑ (Y - M_y_)^2^ = SS_y_ = 2.222

*X and Y Combined*

*N* = 9

∑ (X - M_x_)(Y - M_y_) = 1.722

*R Calculation*

r = ∑ ((X - M_y_)(Y - M_x_)) / √((SS_x_)(SS_y_))

r = 1.722 / √((1.722)(2.222)) = 0.8803

The P-Value is .001735. The result is significant at p < .01.

## Notes

### Competing Interest Statement

The authors have declared no competing interest.

